# Myeloid-cell-specific role of Gasdermin D in promoting lung cancer progression in mice

**DOI:** 10.1101/2022.08.22.504854

**Authors:** C. Alicia Traughber, Gauravi M Deshpande, Kalash Neupane, Mariam R Khan, Megan R McMullen, Shadi Swaidani, Emmanuel Opoku, Santoshi Muppala, Jonathan D Smith, Laura E Nagy, Kailash Gulshan

## Abstract

The activities of the NLRP3 and AIM2 inflammasomes and Gasdermin D (GsdmD), the final executor of inflammasome activity, are implicated in lung cancer pathophysiology but it’s not clear if their contributions promote or retard lung cancer progression. GsdmD plays a role in release of interleukin-1beta (IL-1 β), and the CANTOS trial and recent studies have highlighted a crucial role of IL-1β in promoting lung cancer. Expression of GsdmD was shown to be upregulated in human non-small cell lung cancer (NSCLC) tissue, but its contribution to *in vivo* lung cancer metastasis is not known. Using a metastatic Lewis Lung Carcinoma (LLC) cell model, we show that GsdmD knockout (GsdmD^-/-^) mice form significantly fewer cancer foci in lung, and exhibit markedly decreased lung cancer metastasis. Furthermore, GsdmD^-/-^ mice show a significant ~ 50% increase in median survival rate vs. isogenic WT C57BL6J mice. The cleaved forms of GsdmD and IL-1 β were detected in lung tumor tissue, indicating inflammasome activity in lung tumor microenvironment (TME). Increased migration and growth of LLC cells was observed upon exposure to the conditioned media derived from inflammasome-induced wild type, but not the GsdmD^-/-^, macrophages. Exposure of human A549 lung cancer cells to the conditioned media derived from inflammasome-induced THP-1 macrophages also resulted in increased cell migration. Using bone marrow transplantation, we show the myeloid-specific contribution of GsdmD in lung cancer metastasis. Taken together, our data show that inflammasome activation in macrophages promotes lung cancer growth and migration, and GsdmD plays a myeloid-specific role in lung cancer progression in mice.

## Introduction

Lung cancer accounts for only 13% of all the new cancer diagnoses but 24% of all cancer deaths, making it the leading cause of cancer deaths,, with an estimated 1.8 million deaths (18%), followed by colorectal (9.4%), liver (8.3%), stomach (7.7%), and breast (6.9%) cancers^1, 2^. Epidemiological studies show the role of chronic inflammation in promoting various types of cancers, including lung cancer^3^. Inflammation induced by LPS treatment promotes lung cancer growth in mice^23^. Microbiota, which serves as an endogenous source of LPS, can affect patient responses to cancer immunotherapy^24^ and commensal lung microbiota can modulate levels of Interleukin-1beta (IL-1β) in lung tumor microenvironment^25^. IL-1β is one of the prominent cytokines in the lung tumor microenvironment (TME), promoting tumor invasiveness and metastasis^4–7^. A retrospective study based on the CANTOS trial on human CVD patients^8–10^, showed a 56% reduction in lung cancer incidence in the patients receiving anti-IL-1 β antibody vs. placebo^8^. Activation of NLRP3 or AIM2 inflammasomes allows cleavage of pro IL-1β, followed by release of mature IL-1β in a Gasdermin D (GsdmD) dependent manner^11, 12^, from living as well as pyroptotic cells^12–14^. GsdmD expression was shown to be up-regulated in human non-small cell lung cancer (NSCLC) tissue^15^, while depletion of AIM2 or NLRP3 in lung cancer cells led to reduced growth of cells^16–19^. GsdmD belongs to the family of Gasdermin proteins containing a cytotoxic N-terminal domain and a C-terminal repressor domain shielding the N-terminal domain. After cleavage of GsdmD, the N-terminal fragment gets inserted into the plasma membrane, forming large oligomeric pores, and disrupting membrane integrity^11,12, 20, 21^.

The cleaved N-terminal fragment of GsdmD binds to phosphatidylinositol 4,5-bisphosphate (PIP2) and phosphatidylserine (PS) on the plasma membrane^12, 13^, leading to pore formation, IL-1β release, and pyroptotic cell death ^12–14^. Though studies have implicated inflammasome components in lung cancer^16–19^, but the direct role of GsdmD in lung cancer progression in mouse models or human patients has not yet been explored. We used a syngeneic Lewis Lung Carcinoma (LLC) mouse model, which entails the injection of immunologically compatible cancer cells into fully immunocompetent mice, to determine the role of GsdmD in lung cancer progression. The LLC cell line is highly tumorigenic and is primarily used to model metastasis, being the most used and highly reproducible syngeneic model for lung cancer to date. Furthermore, the LLC model has been successfully used as a preclinical model for evaluating efficacy of anticancer drugs^22^. Here, we tested the role of GsdmD in recipient mice that were injected with LLC on lung cancer progression. We performed bone marrow transplantation (BMT) to determine the myeloid cell specific contribution of GsdmD in promoting LLC tumor progression in mice.

## Results

### Defective IL-1β release in GsdmD^-/-^ mice in polymicrobial cecal slurry induced sepsis model

Previous studies have shown that GsdmD^-/-^ mice are protected against LPS-induced mortality and are defective in IL-1β release upon inflammasome activation ^11,26^. To determine if GsdmD plays a role in the release of IL-1β upon microbial-induced inflammation, we used a polymicrobial cecal slurry injection mouse model^27^. Age and sex-matched C57BL6J-WT and GsdmD^-/-^ mice were injected i.p. with a standardized pool of cryopreserved mouse cecal slurry (4 ml/g body weight). This dose of cecal slurry was selected based on pilot data showing that it did not induce rapid mortality, but still led to a transient decrease in body temperature to 35°C. The IL-1β in plasma at 4h and 6h post-injection increased significantly in WT mice, while the GsdmD^-/-^ mice showed a minimal increase in plasma IL-1β levels *(**Fig. 1***). These data indicate that GsdmD plays a role in microbial-induced IL-1β release in the host.

**Fig.1:**
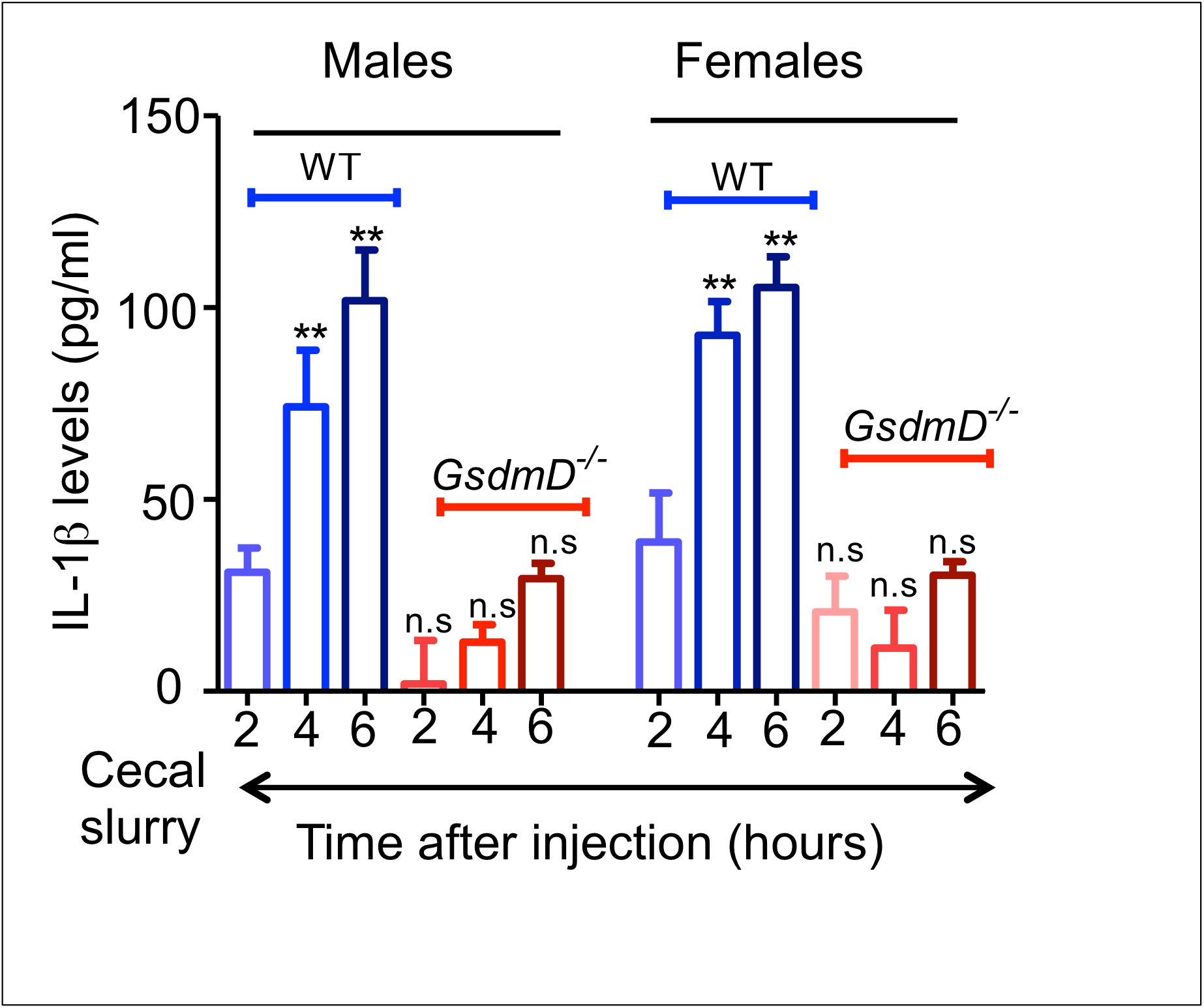
Defective IL-1β release in GsdmD^-/-^ mice upon systemic inflammation. IL-1β ELISA showing levels of plasma IL-1β in WT and GsdmD^-/-^ mice at indicated time periods post cecal slurry injection. Data plotted as mean + SD with N=5 for both males and females, ** show p<0.01 by ANOVA Bonferroni posttest. n.s.= non significant.

### GsdmD^-/-^ mice show markedly reduced LLC metastasis

We tested the effect of GsdmD^-/-^ on lung cancer metastasis in mice by performing i.v. retro-orbital (r.o) injection of LLC cells (2.5 x10^5^ cells per mouse). The heart and lungs were perfused and the visible tumor foci on the lungs were counted. The GsdmD^-/-^ mice showed significantly fewer lung tumor metastatic foci vs. WT mice, with similar effects in both males and females *(**Fig. 2A, Fig. S1A, S1B***). The H & E staining of lung sections from WT mice showed robust cancer growth, while GsdmD^-/-^ mice showed significantly reduced cancer growth *(**Fig. 2B, left panel. Fig. S2A, S2B***). The Masson trichrome staining of WT lung sections showed robust collagen deposition, while GsdmD^-/-^ mice showed significantly reduced collagen deposition ***Fig. 2C, right panel, Fig. S3***). Quantification of the total tumor area and collagen deposition in lung sections showed a marked reduction in % tumor area and collagen deposition in GsdmD^-/-^ mice (***Fig. 2D, 2E***). These data indicate that GsdmD plays a role in lung cancer growth and collagen deposition in cancerous tissue.

**Fig.2:**
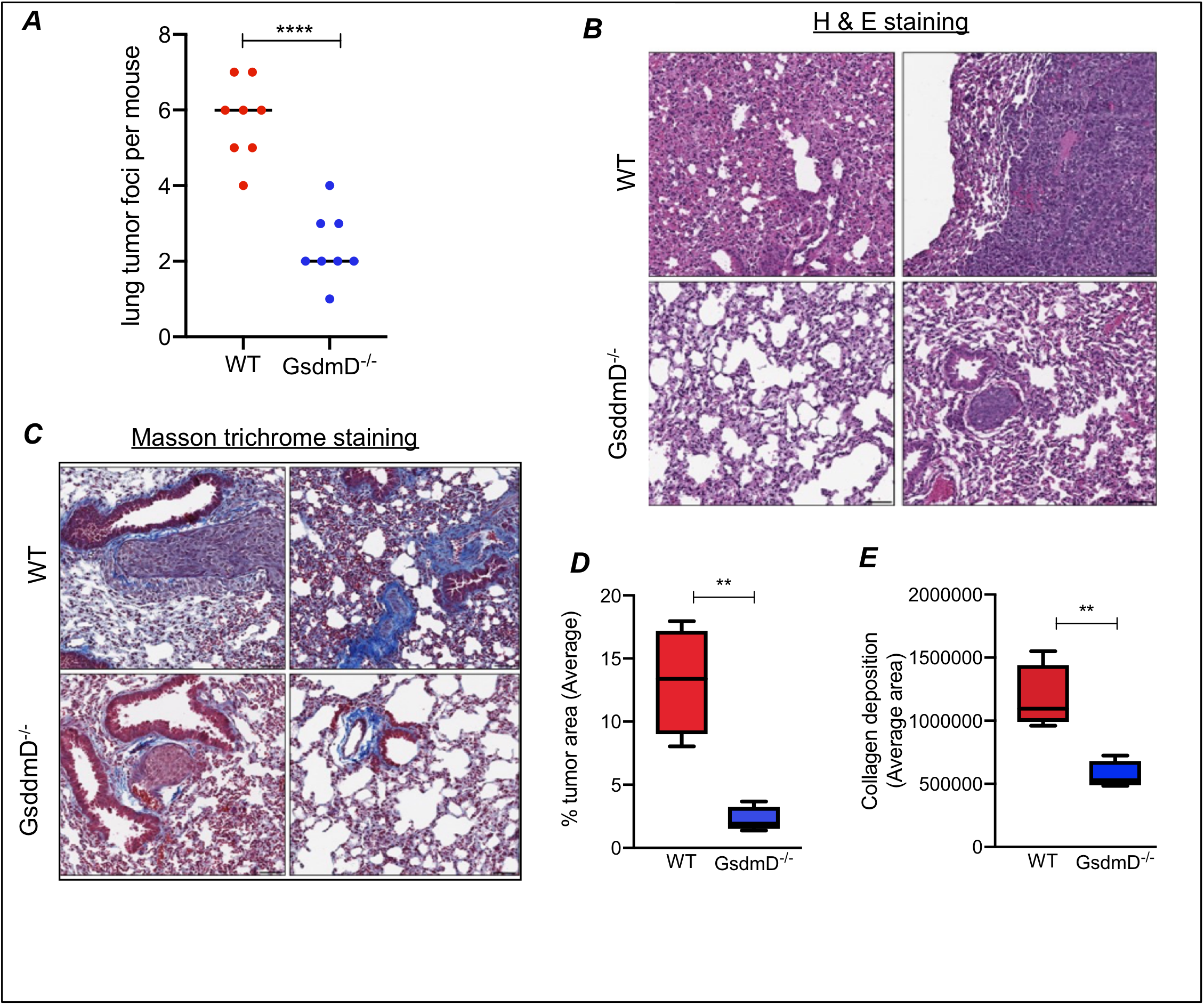
GsdmD^-/-^ mice show reduced lung cancer metastasis. ***A)*** Quantification of metastatic foci in lungs from WT and GsdmD^-/-^ mice (n=8, 4 females, 4 males per strain), ****p=<0.0001, two-tailed t-test. ***B)*** Representative H&E (left panel) staining of lung sections showing tumor growth in WT and GsdmD^-/-^ mice, ***C)*** Representative Masson trichrome staining showing collagen deposition in lung sections from tumor bearing WT and GsdmD^-/-^ mice. ***D)*** Quantification of % tumor area with bar showing average of 4 different sections from WT and GsdmD^-/-^ mice. **p=0.0024, two-tailed t-test ***E)*** Quantification of collagen deposition with bar showing average of different 4 sections WT and GsdmD^-/-^ mice. **p=0.0048, two-tailed t-test

### LLC tumor-bearing GsdmD^-/-^ mice show increased survival

To determine if reduced lung cancer metastasis can lead to increased median survival in GsdmD^-/-^, we performed a Kaplan-Meier survival curve analysis on LLC tumor-bearing WT and GsdmD^-/-^ mice. The mice were injected r.o. with 2.5 x10^5^ LLC cells and were followed till moribund (a marked decrease in body weight, hypothermia, or other conditions requiring immediate euthanasia) or death. The WT mice showed a median survival of ~12 days, while the GsdmD^-/-^ mice showed a significantly higher median survival of ~18 days *(**Fig. 3**).* An independent experiment showed similar results with GsdmD^-/-^ mice showing increased survival days ***(Fig. S4)***. These data indicated that GsdmD promoted lung cancer mortality in mice.

**Fig.3:**
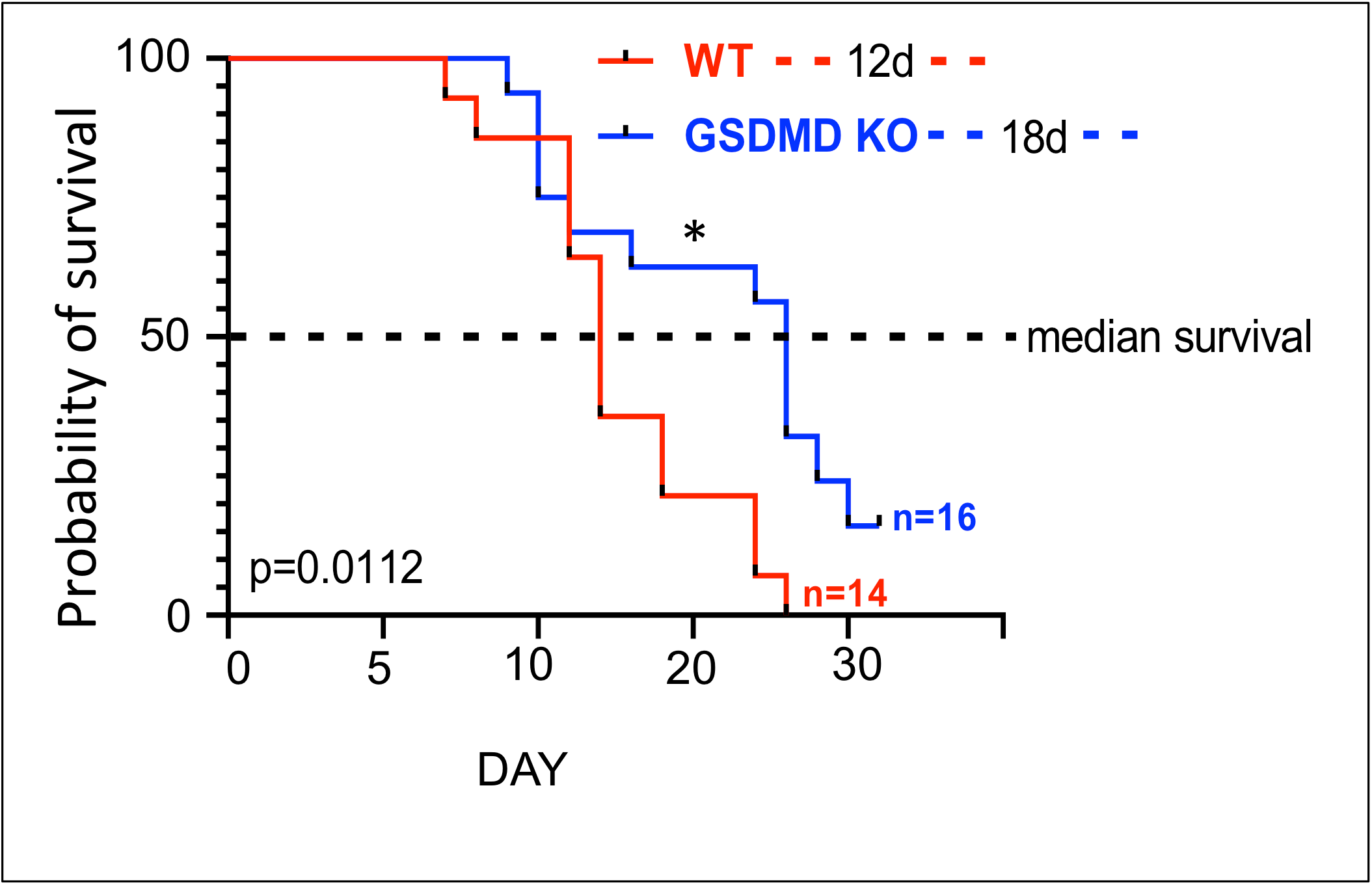
LLC tumor-bearing GsdmD^-/-^ mice show increased survival. Male WT C57BL6J and GsdmD^-/-^ mice were r.o. injected with 2.5 x 10^5^ cells and the survival of mice was determined over period of 25 days. N=14 for WT mice and N=16 for GsdmD^-/-^ mice, p=0.0112 by Log-rank test.

### Inflammasome activity in lung TME

Various studies have indirectly implicated inflammasomes in lung cancer pathophysiology^17,18,28^. To gather evidence for inflammasome activity, the tumor tissue protein extracts were probed with antibodies that recognize the cleaved forms of IL-1β and GsdmD, but not the full-length protein. As a control, adjacent healthy lung tissue was used. As shown in ***Fig. 4***, the cleaved forms of IL-1β and GsdmD were present only in the lysates prepared from tumor tissue, but not from the adjacent healthy lung tissue *(**Fig. 4B**).* These data indicate the presence of functional inflammasome activity in the lung TME.

**Fig.4:**
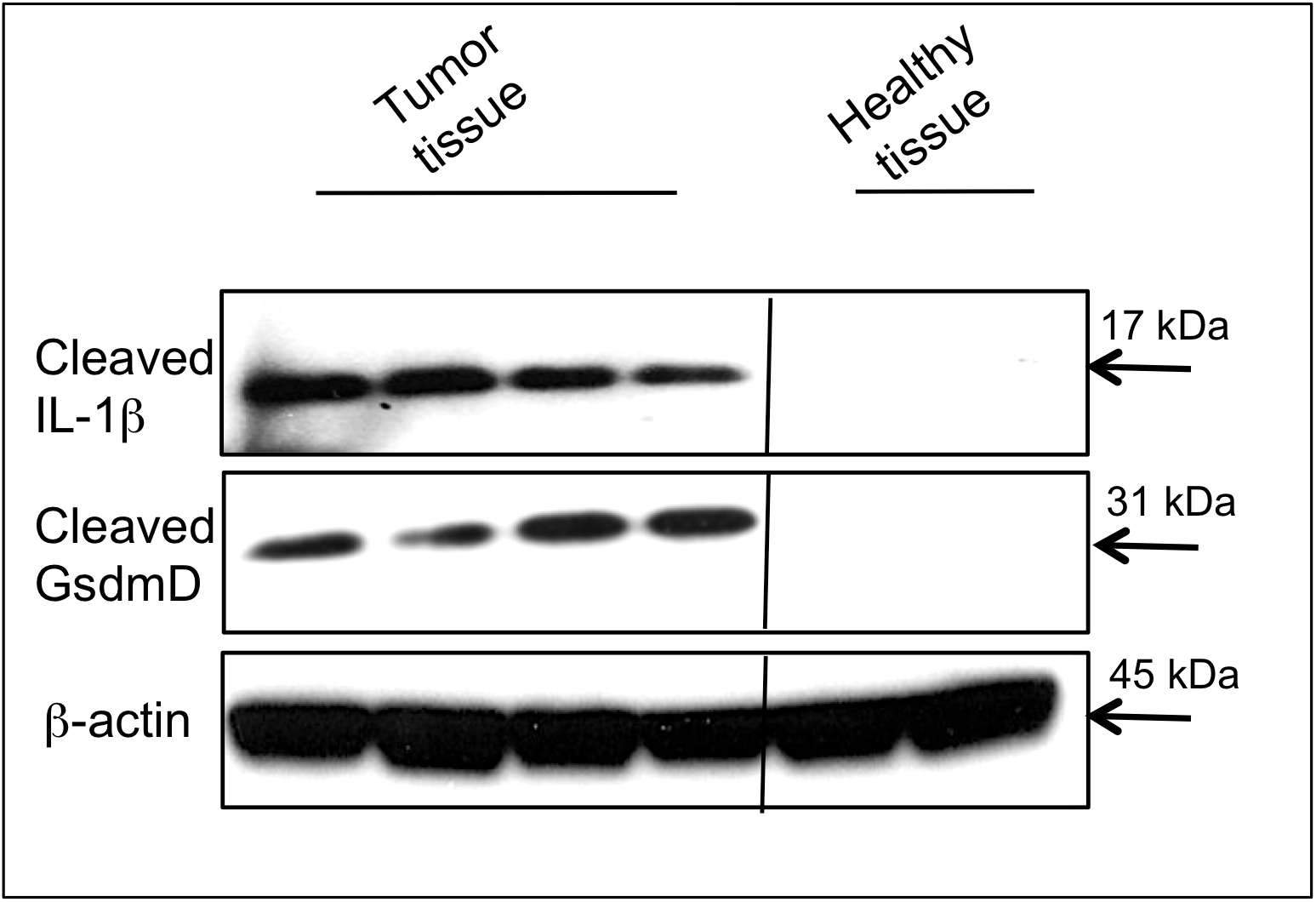
Expression of cleaved GsdmD in lung TME. Western blot analysis of protein extracts from lung tumor tissue and adjacent healthy lung tissue, probed with mouse-specific antibodies against cleaved form of IL-1β and GsdmD. β-actin was used as control.

### Inflammasome-induced macrophages induced cancer cell growth

The growth of lung cancer can be modulated by a variety of factors released in TME by host immune cells as well as by cancer cells. To determine if inflammasome-induced macrophages can modulate cancer cell growth, conditioned media derived from inflammasome-induced WT or GsdmD^-/-^ BMDMs was used. The assembly of NLRP3 or AIM2 inflammasome was induced by treatment with LPS+ATP or with LPS+dA:dT, as described earlier^26, 29^. As shown in ***Fig 5A***, LLC cells incubated with conditioned media from AIM2 inflammasome-induced WT BMDMs induced cell growth by ~ 30%. In contrast, the conditioned media isolated from GsdmD^-/-^ BMDMs failed to show similar robust effects on cell growth. These data indicate that inflammasome assembly in macrophages in lung TME can promote cancer cell growth.

**Fig.5:**
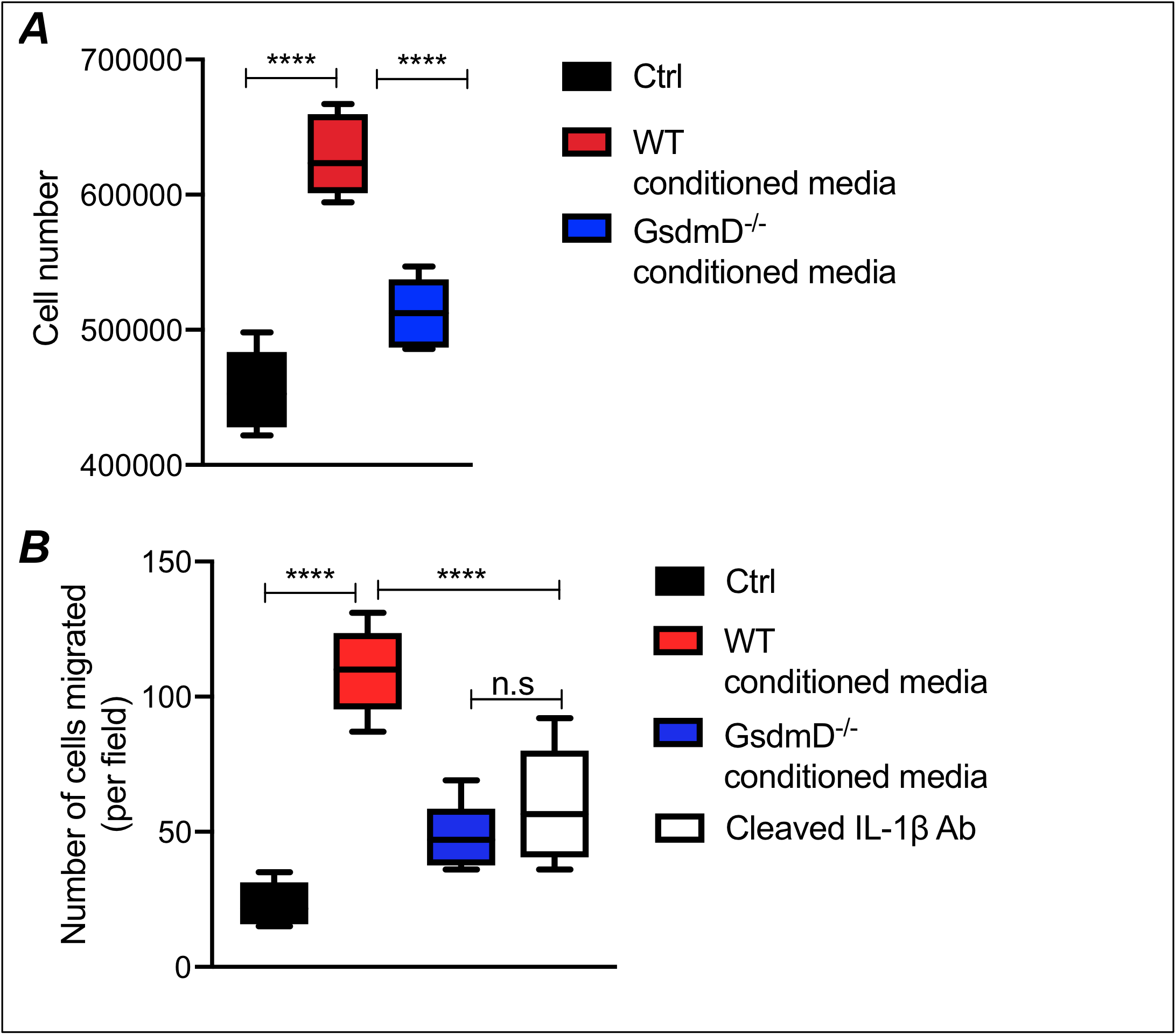
Inflammasome-induced macrophages promoted LLC cell growth and migration. ***A)*** LLC cell growth assay with conditioned media from inflamed WT or GsdmD^-/-^ macrophages, control cells are treated with conditioned media from non-stimulated cells (N=5, whiskers are minimum to maximum, **** represent p<0.0001 for Ctrl vs. WT conditioned media and WT conditioned media vs. GsdmD^-/-^ conditioned media, while Ctrl vs. GsdmD^-/-^ conditioned media showed *, p = 0.0221 with ANOVA using Tukey’s multiple comparisons test. ***B)*** LLC cell trans-well assay migration with conditioned media from inflamed WT or GsdmD^-/-^ macrophages or with WT conditioned media pretreated with antibody against cleaved IL-1β, control cells are treated with conditioned media from non-stimulated cells (N=6, mean ± SD, **** represent p<0.0001 for Ctrl vs. WT conditioned media, **** represent p<0.0001 for WT conditioned media vs. GsdmD^-/-^ conditioned media, and **** represent p<0.0001 for WT conditioned media vs. cleaved IL-1β treatment, while Ctrl vs. cleaved IL-1β treatment showed **, p= 0.0025, Ctrl vs. GsdmD^-/-^ conditioned media showed *, p = 0.0408, and GsdmD^-/-^ conditioned media vs. cleaved IL-1β treatment showed no significant difference (n.s.) with ANOVA using Tukey’s multiple comparisons test.

### Inflammasome-induced macrophages induced cancer cell migration

To determine the mechanism by which inflammasome activation can enhance lung cancer metastasis, the effect of conditioned media derived from inflammasome-induced BMDMs was tested on trans-well migration of LLC cells. The starved LLC were treated with either conditioned media from WT BMDMs or GsdmD^-/-^ BMDMs, and cells were allowed to migrate for 24h. The control cells were left untreated. As shown in ***Fig. 5B***, the conditioned media from inflammasome-induced WT macrophages increased the trans-well migration of LLC cells, while the conditioned media derived from inflammasome-induced GsdmD^-/-^ macrophages failed to show the similar effects. To determine if the increased migration of LLC is indeed due to IL-1β, the conditioned media derived from WT BMDMs was pretreated with antibodies against the cleaved form of IL-1β. As shown in Fig. ***5B***, anti-IL-1β antibody treatment reduced LLC cell migration. These data indicate that the IL-1 β released from the inflamed macrophages induced LLC cell migration. We also tested the effect of conditioned media derived from the inflammasome-activated human THP-1 cells on the migration of human A549 lung cancer cells. The cell migration was determined qualitatively as well as quantitatively via wound healing assay using live-cell video microscopy.

As shown in ***Fig. 6 A and 6B,*** A549 cell migration in the presence of conditioned media derived from inflamed THP-1 macrophages or recombinant human IL-1 β was significantly faster than control cells. Representative video microscopy movies are shown in ***Fig. S5, S6, and S7.*** These data indicated that cytokine release from inflammasome-induced immune cells in TME could induce lung cancer cell migrations and metastasis.

**Fig.6:**
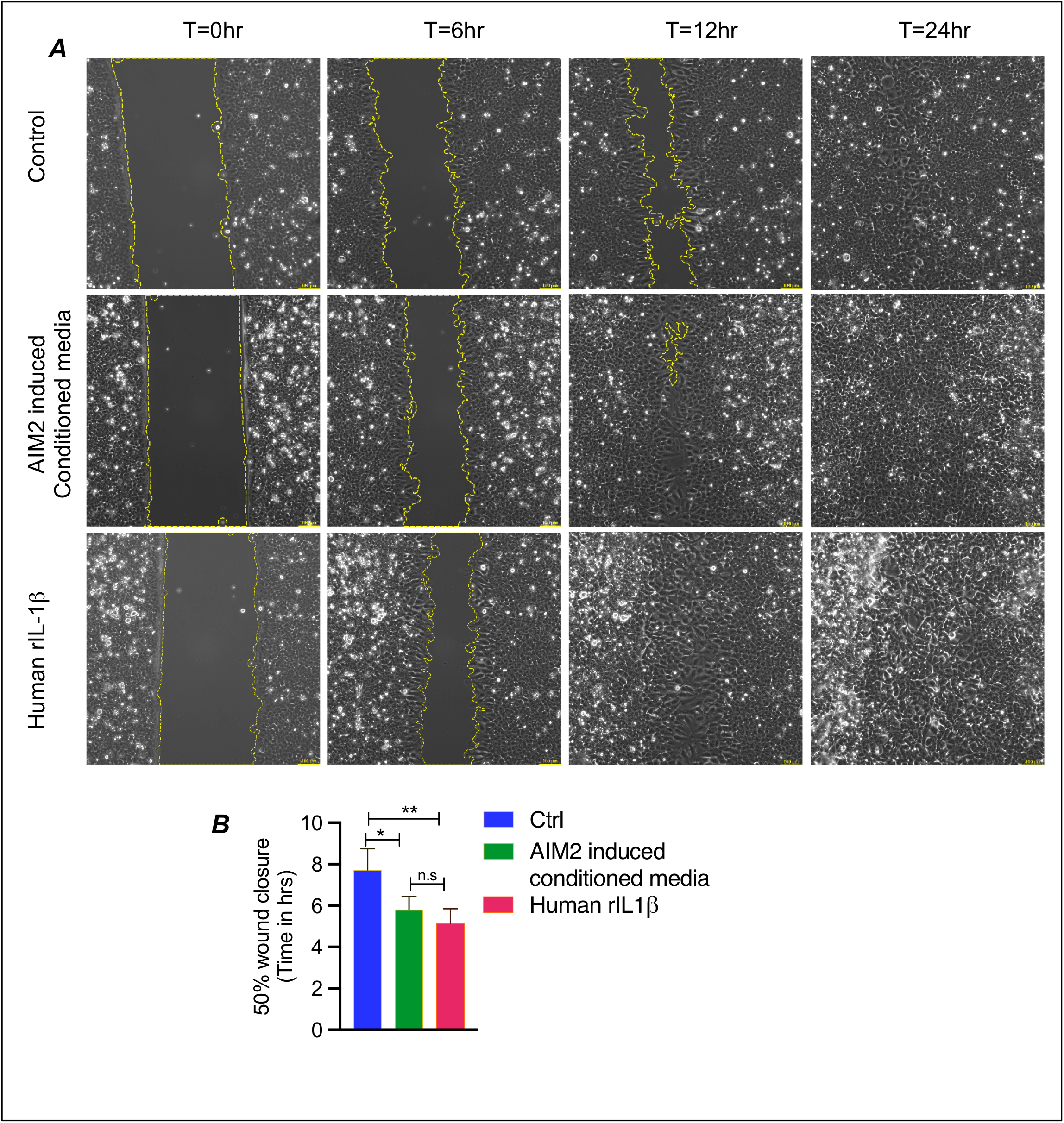
Inflammasome activated THP-1 macrophages promoted migration of human lung cancer cells: ***A)*** Representative images from human A549 cells migration in a wound healing assay with conditioned media from AIM2 inflammasome-induced human THP-1 macrophages or human recombinant IL-1β protein. Control cells are treated with conditioned media from non-stimulated cells. ***B)*** The T_1/2_ (half time) was calculated via live cell video microscopy and plot shows the average of each sample (n=4, mean ± SD, ** represent p=0.0029, * represent p=0.0163, and n.s = non significant, with ANOVA using Tukey’s multiple comparisons test.

### Myeloid cell specific role of GsdmD in lung cancer

To determine if the role of GsdmD in promoting lung cancer is myeloid-cell specific, we performed bone marrow transplant (BMT) assays. A 100% success rate for BMT was achieved, with all the WT mice transplanted with GsdmD^-/-^ bone marrow cells showing the presence of GsdmD^-/-^ myeloid cells in the blood. The BMT strategy and representative successful BMTs are shown in ***Fig. 7A and Fig. 7B***. The 11-12 weeks old male WT mice, transplanted with either WT or GsdmD^-/-^ bone marrow, were given r.o. injection of LLC cells as described above. Mice were followed for survival; endpoints sudden death, moribund, or respiratory distress. As shown in ***Fig. 7C***, the mice transplanted with GsdmD^-/-^ BMT showed a significantly higher survival rate vs. mice transplanted with WT BMT. These data provide the first evidence for the myeloid cell specific role of GsdmD in promoting lung progression.

**Fig.7:**
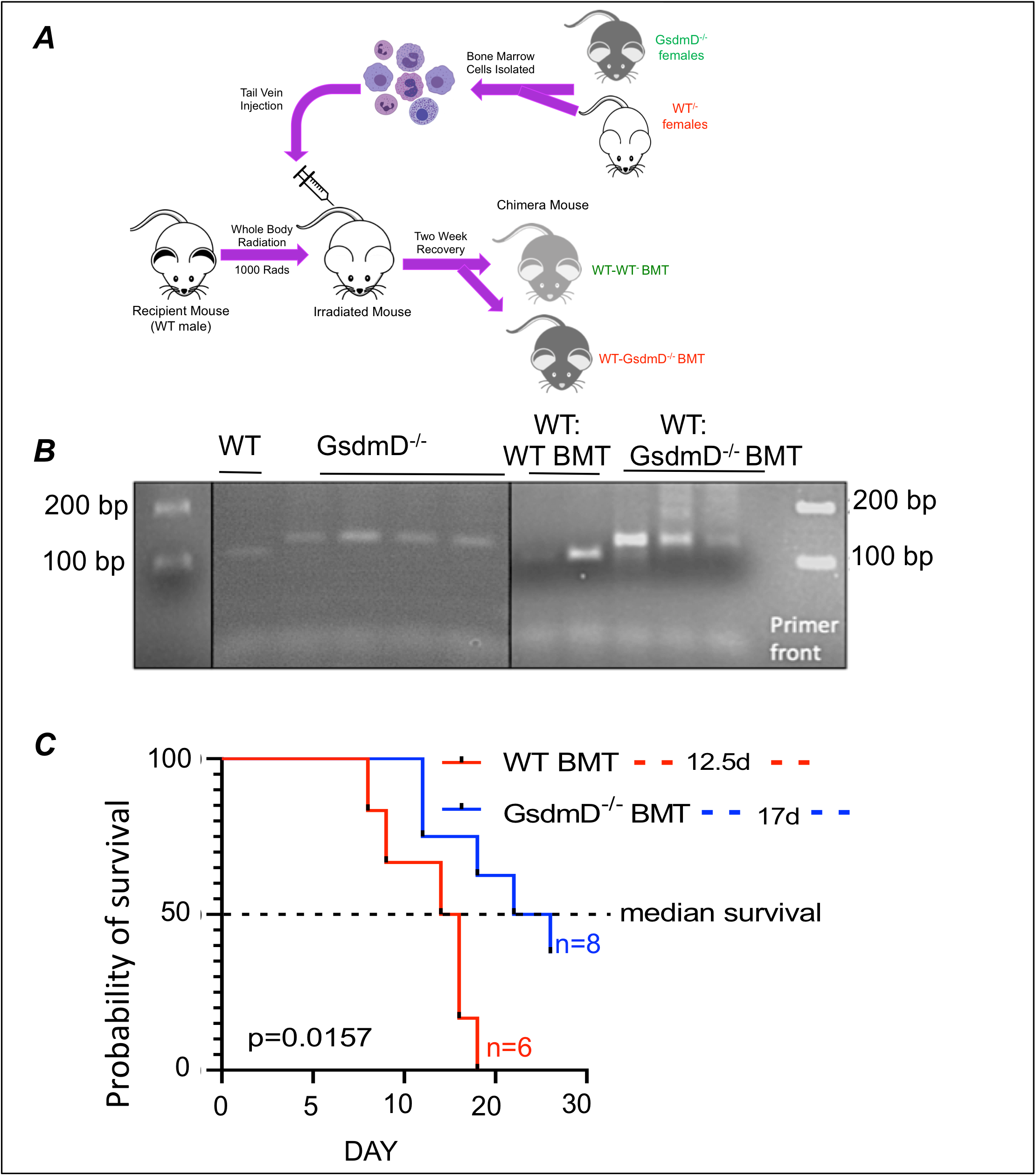
Myeloid-specific role of GsdmD in lung cancer: **A)** Schematic diagram showing BMT strategy. Bone marrow cells were collected from WT or GsdmD^-/-^ females and irradiated WT male mice were injected with bone marrow cells from either WT or GsdmD^-/-^ mice by tail-vein injection. **B)** PCR confirming WT or GsdmD^-/-^ BMT. GsdmD^-/-^ contains 20bp (19bp + 1 bp) insertion with WT sequence “ATGTTGTCAGG” mutated to “ATGcTTGaaggtgtgtggatgcagagTCAGG”. **C)** Survival of WT-WT BMT and WT-GsdmD^-/-^ BMT mice with LLC tumors. N=6 for WT-WT-BMT n=6, and N=8 for WT-GsdmD^-/--^BMT, p=0.0157, by Log-rank test.

### Discussion

Mature IL-1β can be generated by activation of NLRP3 or AIM2 inflammasome. Activation of inflammasomes allows caspase1/11-mediated cleavage of GsdmD in immune cells^11, 12^. The N-terminal fragment of GsdmD is recruited to the plasma membrane, leading to pore formation, IL-1β release, and pyroptotic cell death ^12–14^. Intriguingly, GsdmD is also involved in release of mature IL-1β from living macrophages^30^, indicating role of GsdmD in inflammation is not limited to pyroptosis. Thus, the role of GsdmD from the TME in supporting lung cancer progression may be independent of pyroptosis. The Gasdermin protein family is conserved among vertebrates and is comprised of six members in humans, GsdmA, GsdmB, GsdmC, GsdmD, GsdmE, and DFNB59. Gasdermin proteins are expressed in a variety of cell types including epithelial and immune cells and their role in cancer in diverse, with some members exhibiting tumor promoting properties, while others act to inhibit tumor growth. GsdmE was recently identified as a tumor suppressor in melanoma, breast cancer, and colorectal tumor^31^. GsdmB was shown to promote breast cancer^32,33^, while PD-L1-mediated GsdmC expression was shown to cause a switch from apoptosis to pyroptosis in cancer cells to facilitate tumor necrosis^34^. The role of GsdmD in lung cancer is not consistent across different studies. Multiple studies have shown that depleting the levels of AIM2, NLRP3, or GsdmD in lung cancer cells can inhibit cell growth^16–19^, but the direct role of GsdmD in lung cancer is not fully clear. GsdmD expression was shown to be upregulated in human non-small cell lung cancer (NSCLC) tissue and GsdmD depletion reduced lung cancer growth^15^. In contrast, GsdmD was found to be essential for an optimal cytotoxic T-cells response to cancer cells^35^ indicating the tumor-inhibiting activity of GsdmD. Another elegant study showed the role of IL-1β in lung cancer growth independent of NLRP3 inflammasome, with no effect of GsdmD knockout on lung cancer growth in a skin-graft model^36^. We aimed to determine the *in vivo* role of GsdmD in promoting cancer cell growth in the lungs. Using the metastatic LLC cell model, we found that the knockout of GsdmD in mice markedly reduced LLC cancer progression in lungs *(**Fig. 2, Fig. 3***). Given that the injected LLC cells were not engineered, it is the host GsdmD that supports lung cancer progression. The GsdmD^-/-^ mice showed significantly reduced collagen deposition in cancer tissue vs. WT mice (***Fig. 3***). Collagen in lung TME promotes fibrosis and metastasis of cancer. GsdmD has been proposed to increase collagen production in skin fibrosis^37^, thus GsdmD^-/-^ may be producing less collagen in lung TME, leading to attenuated tumor growth. The GsdmD^-/-^ mice exhibited significantly increased survival upon LLC cell injection vs. WT cells, indicating that GsdmD inhibition may serve as stand-alone or adjuvant therapy to treat lung cancer.

Previous studies implicated inflammasomes in lung cancer cell growth^15, 17, 18, 28^. We found robust activity of inflammasome in lung cancer tissue. The likelihood of inflammasome activity being regulated by immune cells in TME is more plausible rather than lung epithelium being the source of inflammasome activity. Though unlikely, the contribution of host lung epithelial tissue in expression of inflammasome components in cancer tissue cannot be completely ruled out. AIM2 is a receptor that recognizes cytosolic foreign DNA via intracellular Toll-like receptors (TLRs) 7 and 9. One possibility is that DNA from dying cancer cells or DNA fragments from neutrophil extracellular traps (NETosis) are released in lung TME, which are engulfed by macrophages and dendritic cells, allowing activation of AIM2 via TLR7/9.

The presence of cleaved GsdmD in LLC tumor tissue *(**Fig. 4**)* points toward inflammasome activity in lung TME. Though the presence of cleaved GsdmD indicates inflammasome activity, it does not provide foolproof evidence for pyroptosis induced IL-1β in tumor tissue, as living macrophages can also release mature IL-1β^30^. Furthermore, cells can release IL-1β in a GsdmD dependent as well as GsdmD independent manner^14, 38^.

GsdmD is expressed in almost all human organs and tissues including various subsets of leukocytes, while in mice GsdmD is expressed in the gut, colon, urinary tract heart, liver, and lungs. It’s not clear if GsdmD plays a tissue-specific role in promoting lung cancer growth, or which organs are directly involved in GsdmD mediated promotion of lung cancer growth. The function of GsdmD in immune cells is widely studied in the context of bacterial infection but the immune cell specific role of GsdmD in promoting cancer is not known. We found that GsdmD promotes cancer growth in myeloid-cell specific manner *(**Fig. 7**).* GsdmD activation in lung TME can result in an acute rise in IL-1β and other danger signals. High levels of IL-1β in the lung TME can enhance the recruitment of tumor-infiltrating macrophages (TAMs), which are known to promote lung cancer progression and suppress host anti-tumor responses^39^. High density of TAMs in lung cancer correlates with reduced overall patient survival and depletion of TAMs slows lung tumor growth in mice ^40, 41^. We showed the myeloid cell specific role of GsdmD in lung cancer progression, but we cannot entirely rule out the possibility of indirect effects of myeloid GsdmD via modulation of gut or lung microbiota. An elegant study showed that commensal bacteria stimulated MyD88-dependent IL-1β and IL-23 production from myeloid cells^25^. There is a possibility of lung microbiota being affected in WT mice with GsdmD^-/-^ BMT vs, WT mice with WT BMT. GsdmD, due to its bactericidal activity, may tilt the lung microbiota population toward a cancer-promoting rather than healthy microbiota profile.

Targeting GsdmD along with IL-1β may serve as a better adjuvant therapy than targeting IL-1β alone, as GsdmD blockage can prevent not only the IL-1β release, but can also prevent highly inflammatory pyroptotic cell death, formation of neutrophils extracellular traps, and leakage of other pro-inflammatory cytokines and cellular material in the lung TME. GsdmD inhibition may also reduce lung cancer by inducing apoptosis in inflamed TAMs. Recent studies from our group and others have shown that GsdmD^-/-^ macrophages are more susceptible to apoptosis under inflammatory conditions, such as overload of cholesterol^26, 42^. Thus, the inflammatory environment of lung TME may promote apoptosis in GsdmD^-/-^-TAMs vs. highly inflammatory pyroptotic cell death in WT-TAMs.

The mechanistic details of inflammasome-GsdmD-IL-1 β nexus in promoting lung cancer are still not fully clear. The cross-talk between GsdmD activity in macrophages, neutrophils, and other immune cells such as Tregs can modulate the host response to cancerous growth in lungs, but delineating the intricate details of the regulatory network involved in this process requires further studies.

## Acknowledgments

This research was supported by Case Comprehensive Cancer Center pilot grant and Cleveland State University startup, NIH-NHLBI R01-158148 grant to K.G.

## Disclosures

Authors declare no financial or non-financial competing conflict of interests.

## Material and Methods

### Cell lines and BMDMs

THP-1 cells were cultured in RPMI medium supplemented with 10 % FBS, penicillin–streptomycin antibiotics, and 0.05 mM 2-mercaptoethanol. THP-1 cells were differentiated into macrophages using 100ng/ml phorbol 12-myristate 13-acetate (Sigma P8139) for 3 days. Bone marrow-derived macrophages (BMDMs) were generated from bone marrow cells of mice. WT or GsdmD^-/-^ mice were euthanized by CO_2_ inhalation and femoral bones were removed. The marrow was flushed out of the bones into a 50 ml sterile tube using a 10 ml syringe with a 26-gauge needle filled with sterile DMEM. Cells were centrifuged for 5 min at 1,800 rpm at 4°C, followed by two washes with sterile PBS. The bone marrow cells were suspended in sterile-filtered BMDM growth media (DMEM with 7.6% fetal bovine serum, 15% L-cell conditioned media, and 0.76% penicillin/streptomycin mixture) and plated in culture dishes and incubated at 37°C for 14 days. Cell media was replaced every 2–3 days for 2 weeks. The cells were routinely visualized under microscope for proliferation and differentiation into confluent BMDMs.

### Mice maintenance/diet

All animal experiments were pre-approved by the Cleveland Clinic IACUC and the Cleveland State University IACUC. WT C57BL6J were purchased from The Jackson Laboratory and C57BL6J-GsdmD^-/-^ mice were described before^26, 43^. Mice were maintained in a temperature-controlled facility with a 12-h light/dark cycle with free access to food and water. The standard chow diet (SD, 20% kcal protein, 70% kcal carbohydrate and 10% kcal fat, Harlan Teklad) was used for regular maintenance and breeding.

### LLC metastasis R.O injection model

LLC cells were cultured in regular growth media and 2.5×10^5 cells were injected into each mice. Male WT or GsdmD^-/-^ mice (10-11 weeks of age) were put under isoflurane anesthesia and given i.v. injection into the retroorbital plexus in a volume of 0.25 ml. For injections, the skin above the eye was drawn back to protrude the eye slightly and 28-gauge needle (not to exceed 1/2” to avoid trauma) was inserted at an angle of ~45 degree, through the inferior fornix conjunctiva membrane. The needle was gently removed to prevent injury to the eye. Eyelid was closed and mild pressure was applied to the injection site with a gauze sponge.

### Cell migration assay

LLC cells were treated with conditioned media from inflammasome-induced macrophages as indicated and 5 × 10^4^ cells per well were seeded in top chambers of the transwell chamber (8 μM pore, 24-well plate, Millipore-Sigma) in FBS-free media with membrane inserts. The lower compartment contained DMEM supplemented with 10% FBS as attractant. The cells were incubated at 37°C for 24h. The cells on the lower side of the insert membrane were fixed with 5% glutaraldehyde for 10 min, followed by staining with 1% crystal violet in 2% ethanol for 30 min. The inserts were washed extensively with PBS to remove excess dye. The cells in the upper compartment of the insert were gently removed by gently wiping with a cotton swab. The insert was completely dried and the number of cells on the lower side of the filter were counted under a microscope.

### Wound healing assay

Human A549 lung cancer cells were cultured and treated with vehicle, conditioned media from inflammasome-induced macrophages, or human recombinant IL-1β (rIL-1β) as indicated. Images were acquired at 10X/0.40NA, every 15 minutes for 24 hours using a Leica DMI6000 inverted microscope (Leica Microsystems, GmbH, Wetzlar, Germany) equipped with a Hamamatsu Orca Flash4 camera (Hamamatsu Photonics, Shizuoka, Japan) For live cell video microscopy, the system *was* maintained at 37°C with 5% CO2 throughout the experiment. Image analysis was done using *I*mage-Pro Plus 10 (Media Cybernetics, Inc., Rockville, Maryland, USA) to quantify the time it takes for the gap to close to the 50% of its original area.

### Trichrome/H&E staining and quantification

Paraffin processing, embedding and sectioning of the formalin fixed lung tissue was performed at Cleveland Clinic histology core. Lung sections were stained with Hematoxylin and Eosin (H&E) and Masson’s Trichrome dyes. Leica Aperio AT2 slide scanner (Leica Microsystems, GmbH, Wetzlar, Germany) was used to scan whole slides at 20X magnification to get a resolution of 0.5micron/pixel. The images were converted to tiff files using Aperio ImageScope software. Tumor areas in WT and GsdmD^-/-^ mice were manually chosen and quantified on H&E stained slides using Image-Pro Plus 10 (Media Cybernetics, Inc., Rockville, Maryland, USA) Percent tumor area was calculated as followed: Percent tumor area = (Total tumor area ÷ Total tissue area)×100. Collagen content in WT and GsdmD was quantified on Trichrome stained slides using Image-Pro Plus 10 (Media Cybernetics, Inc., Rockville, Maryland, USA)

### Western blotting

The excised tumors were weighed and equal size tissue was homogenized in tissue lysis buffer (TPER, Thermo Scientific, 78510) supplemented with EDTA free protease inhibitor cocktail (Thermo Scientific, 87785), with ratio of ~0.1g of tissue to 1 mL T-PER lysis buffer. The cell debris was separated by centrifugation at 10,000g for 10 min. After discarding the pellet, the protein concentration was determined using the BCA protein assay (Pierce). 10-50 μg of cell protein samples were resolved on Novex 4-20% Tris-Glycine Gels (Invitrogen) and transferred onto polyvinylidene fluoride membranes (Invitrogen). Blots were incubated with 1:1000 rabbit polyclonal antibody against cleaved IL-1β (Cell Signaling #63124), cleaved GsdmD (Cell Signaling #50928), or β-actin (Cell Signaling #8457 for 4h at RT or for 16h at 4° C. The membranes were washed with PBST for 5 min x 3 times and incubated with 1:15,000 horseradish peroxidase-conjugated goat anti-rabbit secondary antibody (Biorad) for 2h. The signal was detected with an enhanced chemiluminescent substrate (Pierce) and membranes were imaged using iBright™ CL750 Imaging System (Life Technologies, A44116). The western blot bands were quantified via densitometry using the iBright Analysis Software.

### Generation of myeloid-specific KO of GsdmD in mice

To isolate bone marrow, donor mice (female, 6-12 weeks of age) were sacrificed by CO2 asphyxiation followed by cervical dislocation. Femurs and tibias were dissected free of tissue and flushed with sterile PBS. Each mice yields approximately 4 x10^7^ cells, thus each donor can provide bone marrow for ~3 recipients. Bone marrow cells were centrifuged and suspended in sterile PBS prior to injection. Recipient mice: 5-week-old female or male mice were lethally irradiated (600RAD 2x from the cesium source of the Shepard Irradiator). Four hours later, mice received ~ 10^7^ bone marrow cells in 200 μl sterile PBS by intravenous injection into the tail vein. Mice were monitored daily for signs of sepsis (lack of eating, drinking, socializing, grooming, hunched posture, shivering, lethargy). Four weeks after irradiation, mice were treated with a single dose of liposomal clodronate to deplete sessile Kupffer cells that are radio-resistant. Kupffer cells are replenished within 7 days after clodronate. Clodronate containing liposomes were diluted in sterile saline and injected in a 0.20 ml volume of a 1mg/ml solution and Clodronate was given via tail vein injection.

### Statistical Analysis

Comparisons of 2 groups were performed by a 2-tailed t test, and comparisons of 3 or more groups were performed by ANOVA with Bonferroni posttest. All statistics were performed using Prism software (GraphPad). For survival studies, the log-rank test, non-parametric test, was used.

## Notes

### Competing Interest Statement

The authors have declared no competing interest.

