## supplementary material for "Myeloid-cell-specific role of Gasdermin D in promoting lung cancer progression in mice"

Running title: GsdmD promotes lung cancer.

C. Alicia Traughber[1,2,3], Gauravi M Deshpande[4#], Kalash Neupane[1,2#], Mariam R Khan[1,2#], Megan McMullen[5,] Shadi Swaidani[3] , Emmanuel Opoku[3], Santoshi Muppala[3], Jonathan D Smith[3], Laura Nagy[5,6], and Kailash Gulshan[1,2,3]*.

### *Supplementary figures*

***Fig. S1: A)*** Representative image showing LLC tumor foci on lungs of mice injected with LLC cells***. B)*** WT vs. GsdmD$^{-/-}$ lungs. Tumor foci are marked with black arrows.

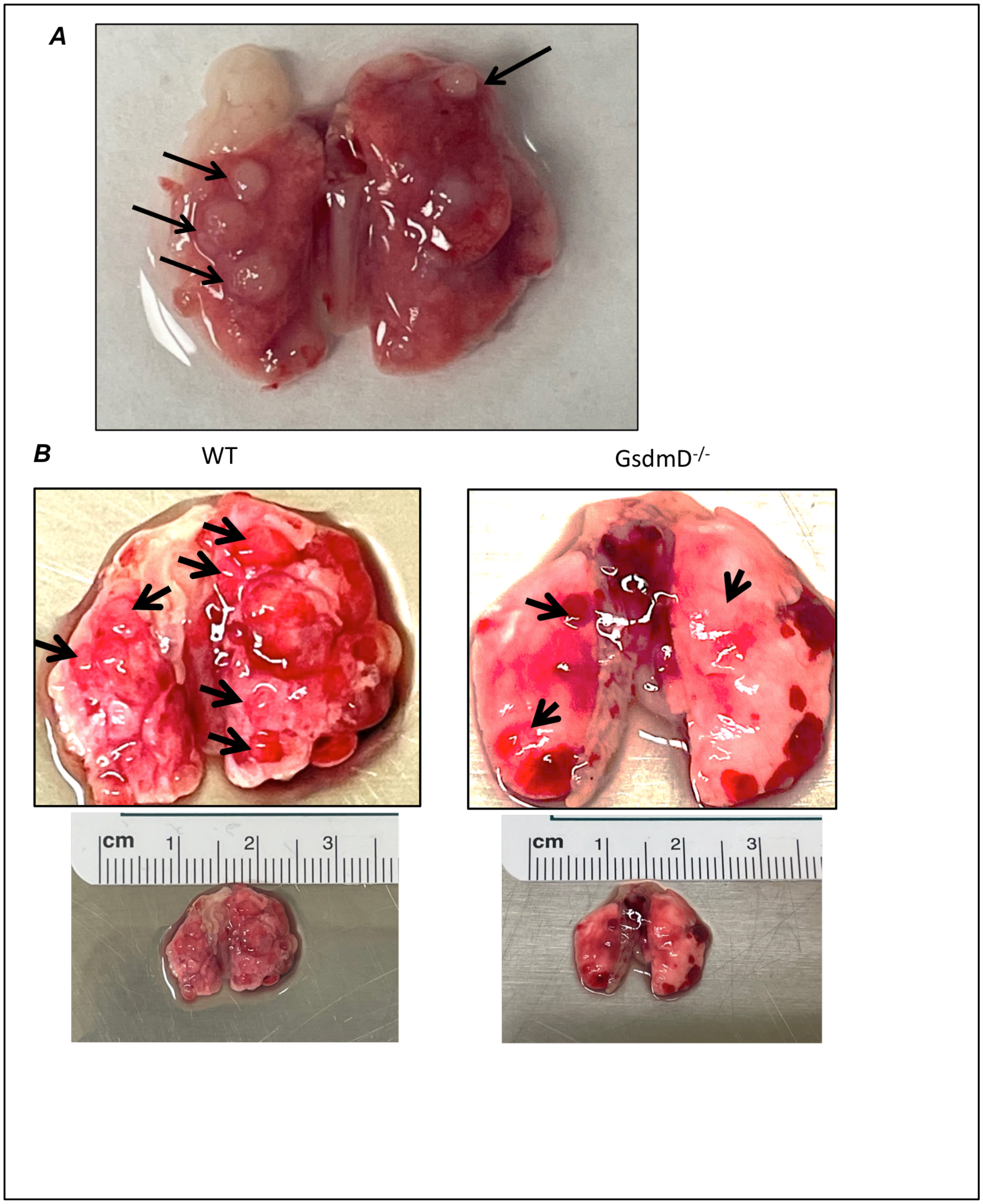

***Figure S2:*** **A) F**ull image of H & E staining sections from WT and Gsdmd$^{-/-}$ mice**. B**) H & E staining images from various **l**ung tissue sections from tumor bearing WT and GsdmD$^{-/-}$ mice.

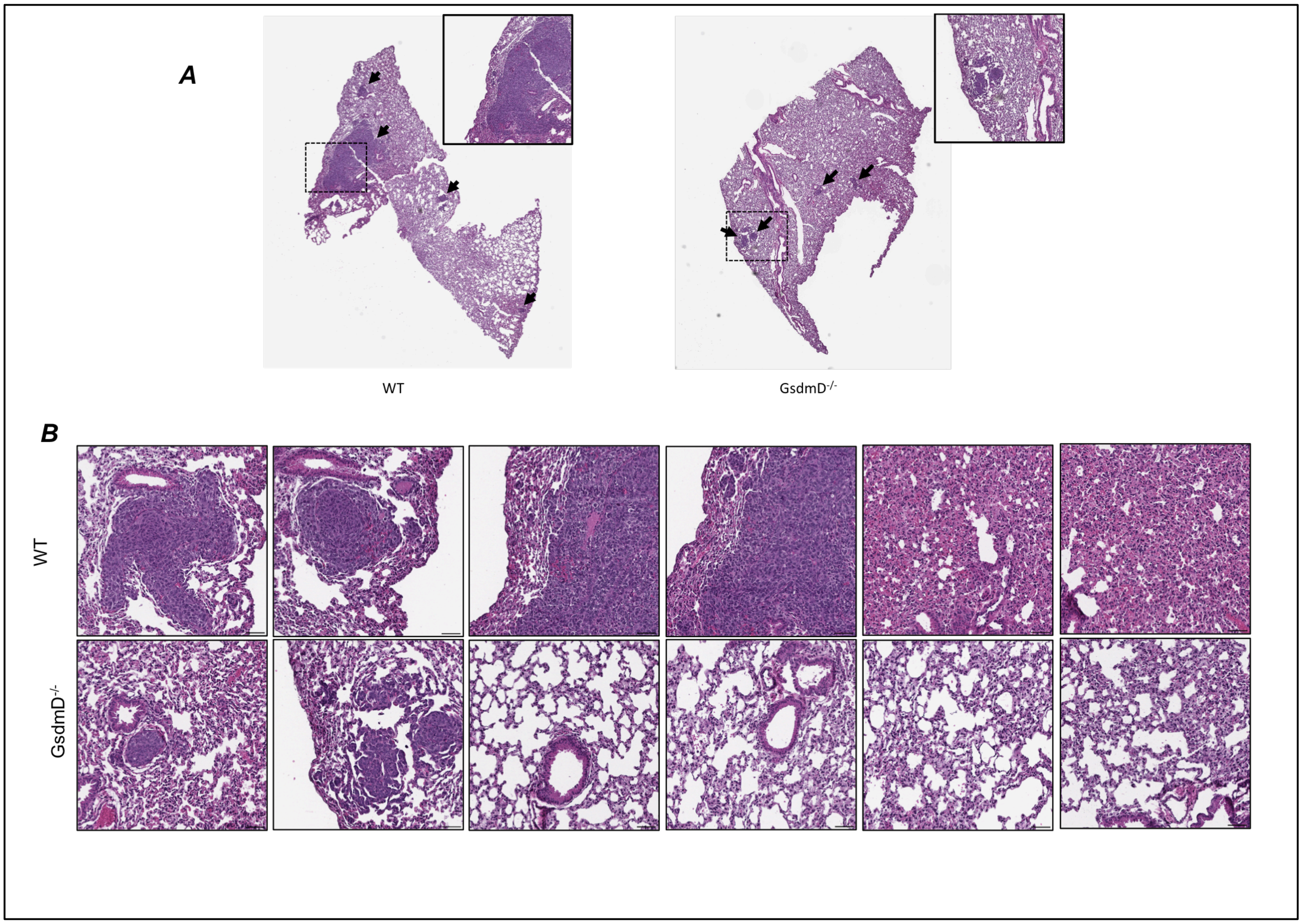

***Figure S3:*** **A) F**ull image of Trichrome staining sections from WT and Gsdmd$^{-/-}$ mice GsdmD$^{-/-}$ mice.

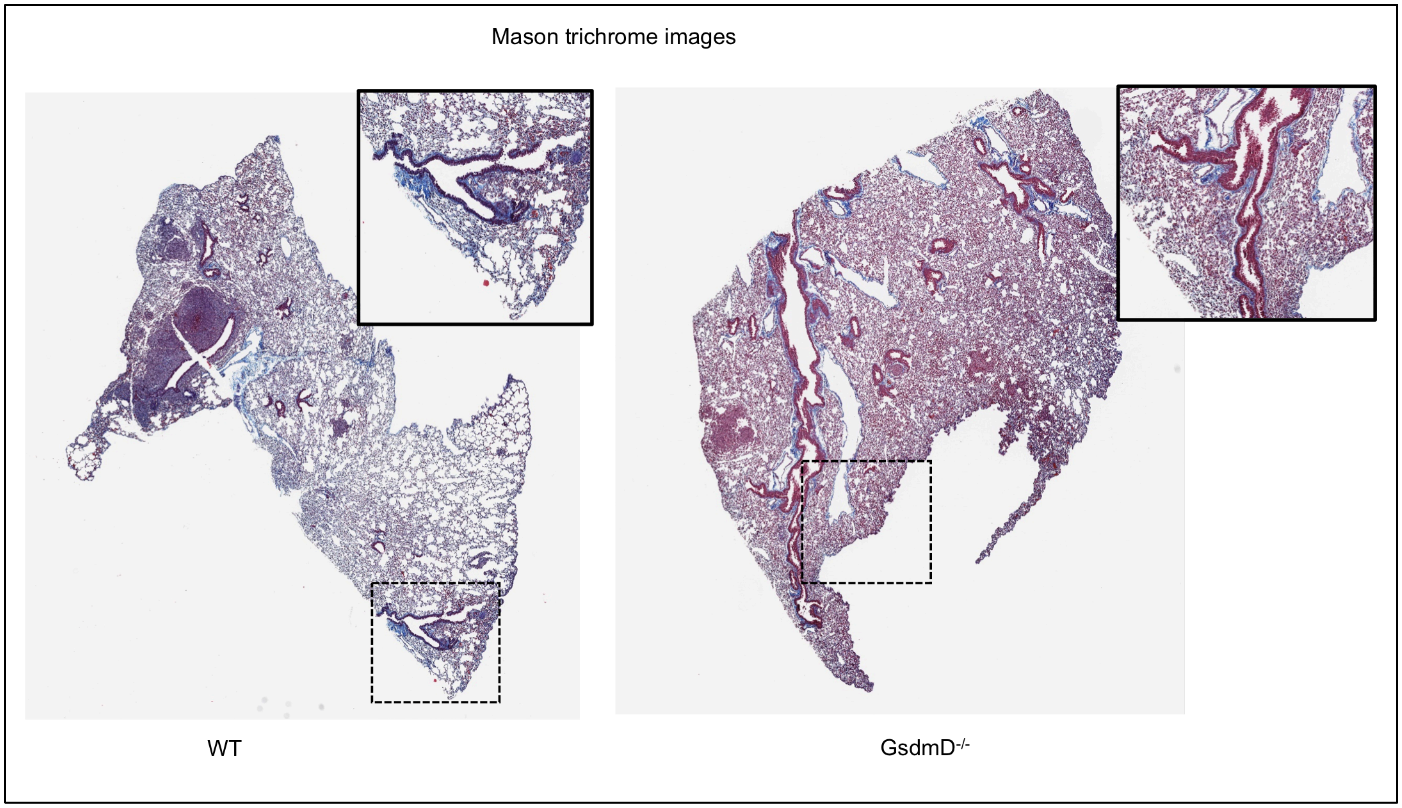

**Fig.S4: LLC tumor-bearing GsdmD$^{-/-}$ mice show increased survival.** Male WT C57BL6J and GsdmD$^{-/-}$ mice were r.o. injected with LLC cells and the survival of mice was determined over period of 25 days. N=4 for WT mice and N=6 for GsdmD$^{-/-}$ mice, p=0.031 by Log-rank test.

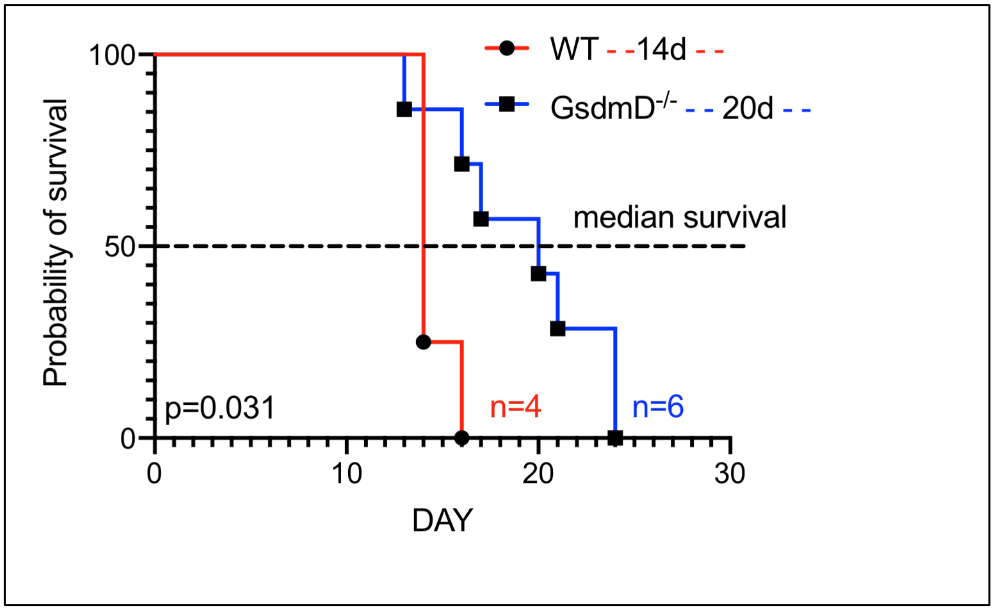

***Figure S5:*** Video-microscopy showing migration of untreated human A549 cells.

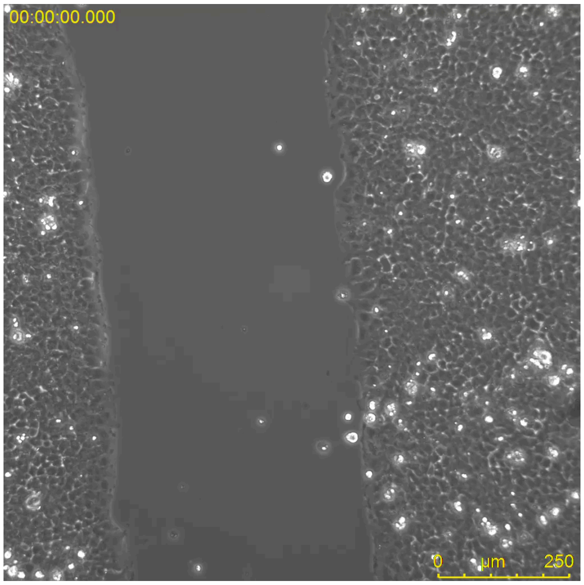


***Figure S6:*** Video-microscopy showing migration of human A549 cells treated with conditioned derived from THP-1 macrophages.

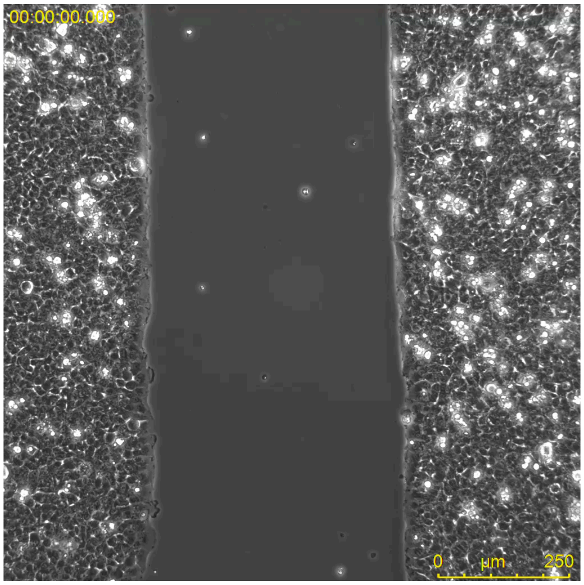

***Figure S7:*** Video-microscopy showing migration of human A549 cells treated with recombinant human IL-1β.

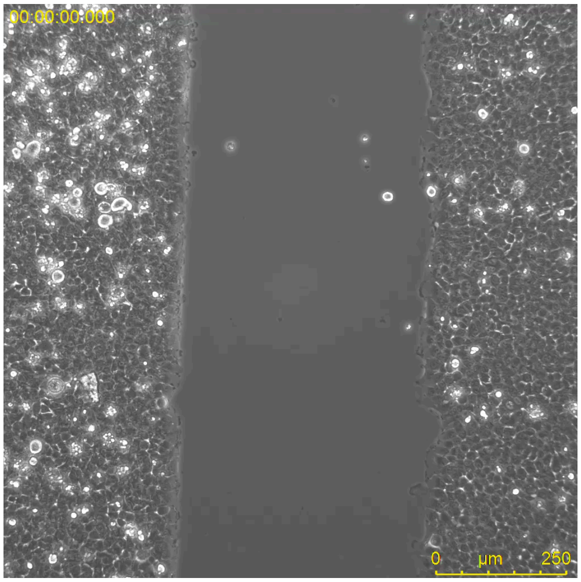
